# A *de novo* transcription-dependent TAD boundary underpins critical multiway interactions during antibody class switch recombination

**DOI:** 10.1101/2022.04.26.489407

**Authors:** Julia Costea, Ursula E. Schoeberl, Daniel Malzl, Maximilian von der Linde, Johanna Fitz, Marina Makharova, Anton Goloborodko, Rushad Pavri

## Abstract

Conflicts between transcription and cohesin-mediated loop extrusion can majorly influence 3D chromatin architecture but whether these structural changes affect biological function is unknown. Here, we show that a critical step in antibody class switch recombination (CSR) in activated B cells, namely, the juxtaposition (synapsis) of donor and acceptor switch (S) recombination sequences at the immunoglobulin heavy chain locus (*Igh*), occurs at the interface of a *de novo* topologically associating domain (TAD) boundary formed via transcriptional activity at acceptor S regions. Using Tri-C to capture higher-order multiway chromatin conformations, we find that synapsis occurs predominantly in the proximity of distal 3’ CTCF-binding sites and that this multiway conformation is abolished upon downregulation of transcription and loss of the TAD boundary at the acceptor S region. Thus, an insulating *de novo* TAD boundary created by the conflict between transcription and loop extrusion plays a direct role in the mechanism of CSR.

## INTRODUCTION

The relationship between chromatin structure and function is fundamental to our understanding of genome organization and gene regulation. Transcription can create topological barriers for chromatin loop extrusion by cohesin complexes. As a result, conflicts between transcription and loop extrusion can majorly influence chromatin structure at active genes and enhancers (*1–6*). However, depletion of cohesin or the major loop-anchoring architectural protein, CTCF, has surprisingly minor effects on global transcription (*3, 7–11*). Thus, whether the structural changes caused by conflicts between loop extrusion and transcription contribute directly to biological function remains an important question in genome and cell biology.

One physiological pathway where loop extrusion and transcription have been proposed to work in concert is antibody class switch recombination (CSR) in B lymphocytes (*12–15*). CSR generates the various antibody isotypes that mediate key effector functions during the humoral immune response (*16*). CSR occurs at the *Igh* locus in mice and involves the exchange of the C*μ* constant region of the constitutively active *Ighm* constant region gene for one of six additional downstream constant (C) genes (*Ighg3, Ighg1, Ighg2b, Ighg2a, Ighe* and *Igha*) spread over 200 kb (Fig. 1A) (*17*). The constant region genes are followed by the 3’ regulatory region (3’RR) which consists of four enhancers termed hypersensitive sites (hs) 3a, 1,2, 3a and 4 (hs3a, hs1,2, hs3b and hs4) (Fig. 1A) (*18, 19*). The 3’ end of the locus is marked by ten CTCF-bound insulator sites termed the 3’ CTCF binding element (3’CBE) (Fig. 1A). Each *Igh* constant region gene consists of a germline I promoter, a repetitive switch (S) sequence and the C exons (Fig. 1A). B cell activation results in germline non-coding transcription from one or more downstream *Igh* constant region genes (*Ighg1* in Fig. 1A) and the expression of activation-induced cytidine deaminase (AID) (*20, 21*). The mutagenic activity of AID initiates the CSR reaction cascade leading to DNA breaks in donor (S*μ*) and any transcribed downstream acceptor S region (S*γ*1 in Fig. 1A) (*22, 23*). The DNA breaks are joined via deletional recombination resulting in CSR and the expression of a new antibody isotype (IgG1 in Fig. 1A) (*24*).

**Figure 1:**
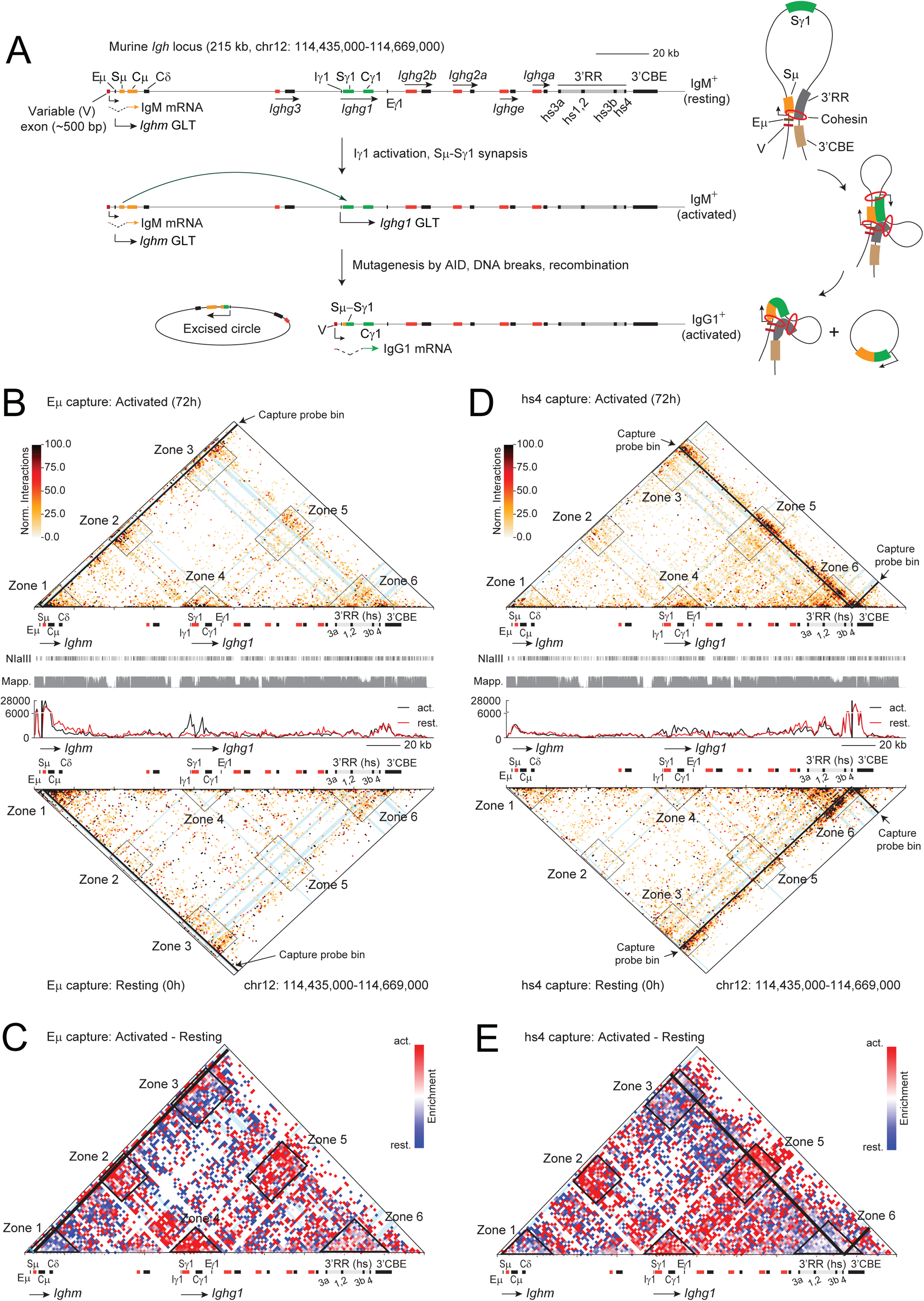
Multi-way interactions of Eμ and hs4 enhancers in resting and CSR-activated B cells. A. Overview of IgG1 CSR at the murine *Igh* locus. The locus is drawn to scale. The *Igh* genes are indicated as well as the Eμ, Eγ1 and 3’RR enhancers and the 3’CBE which contains ten CTCF-binding sites. Each *Igh* gene consists of a promoter (only the Iμ and Iγ1 promoters are shown), repetitive switch (S) sequences (orange for Sμ, green for Sγ1 and red for all others) and constant (C) region exons (orange for Cμ, green for Cγ1 and black for all others). The four enhancers (hypersensitive sites) comprising the 3’RR are indicated as black bars within the 3’RR (hs3a, hs1,2, hs3b and hs4). In resting, IgM-expressing (IgM+) B cells, the variable region (V) exon is spliced to the first constant (C) region (Cμ) resulting in IgM mRNA expression. A non-coding germline transcript (GLT) is also made from the Iμ promoter (not indicated) that lies at the 3’ end of the Eμ enhancer. In the present study, activation of B cells leads to germline transcription predominantly from the Iγ1 promoter resulting in the *Ighg1* GLT. B cell activation also triggers expression of AID which generates lesions in donor (Sμ) and acceptor switch (Sγ1) regions. Processing of these lesion by DNA repair proteins leads to DNA double-strand breaks in the S regions that are used as substrates for deletional recombination between Sμ and Sγ1 via non-homologous end-joining. This results in the excision of the intervening DNA as a circular product and the generation of IgG1 mRNA and IgG1-expressing (IgG1+) B cells. The chromatin loop diagram on the right depicts the current model of CSR wherein cohesin-mediated loop extrusion creates a 3-way conformation that aligns active S regions in the proximity of the 3’RR. This results in the proposed coupling of transcriptional activation and S-S synapsis (Wuerffel et al. Immunity 2007, Zhang et al. Nature 2019).
B. Tri-C analysis from resting (0h, bottom matrix) and activated (72h, top matrix) primary B cells using Eμ as the capture viewpoint. The matrices consist of 1 kb bins spanning the *Igh* locus. Displayed are the reporter fragment counts from ≥3-way interaction fragments normalized by the number of NlaIII restriction sites per bin and adjusted to a total of 300,000 counts per matrix. The bins containing the Eμ capture probe asre blanked (black stripes). The annotation of genes, enhancers and 3‘CBE (elaborated in Fig. S1A) is shown with switch (S) regions of each *Igh* gene indicated as a red bar. The blue lines within the matrix indicate the location of bins corresponding to Eμ enhancer, Iγ1 promoter, Eγ1 enhancer and the four enhancers (hs sites) within the 3’RR. The histogram in between the matrices shows the cumulative contact frequency in each bin of the matrix. A mappability track in grey (Mapp.) is also provided. Note that Sγ1 is poorly mappable due to its highly repetitive sequence as is the core of Sμ and a portion of the palindromic sequence between hs1,2 and hs3b. The six zones of contact enrichment are encircled and labeled (zones 1-6). Zone 1: Eμ with the *Ighm* locale. Zone 2: Eμ-*Ighm*-*Ighg1*. Zone 3: Eμ-*Ighm*-3’RR-3’CBE. Zone 4: Eμ-*Ighg1.* Zone 5: Eμ-*Ighg1*-3’RR-3’CBE. Zone 6: Eμ-3’RR-3’CBE. Zones 5 and 6 represent biological 3-way interactions.
C. A difference matrix (activated - resting) obtained from the data in B. Colored bins show enriched contacts in activated cells (red) or resting cells (blue). The same zones (1-6) are highlighted as in the matrix in B.
D. Tri-C analysis as in B but from the hs4 viewpoint, the last enhancer in the 3’RR. The hs4-containing bins are blanked. Zones 1-6 are the same as in B above with zone 2 representing biological 3-way interactions (hs4-*Ighm*-*Ighg1*).
E. Difference matrix as in C (activated - resting) obtained from the data in D and described in C above.

Importantly, the events critical for CSR, namely, activation of transcription by the 3’RR, as well as the juxtaposition of donor and acceptor S regions (S-S synapsis) essential for bringing DNA breaks into proximity, require long-range interactions involving the formation of 50-200 kb chromatin loops via cohesin-mediated chromatin loop extrusion (*13, 25*). In resting B cell, a basal loop is formed via interactions between the E*μ* and the 3’RR enhancers. Upon activation, interactions between donor and newly transcribed acceptor S regions as well as interactions between the 3’RR and acceptor S regions are observed(*12, 25–28*). This has led to a model wherein the recruitment of an acceptor I promoter (I*γ*1 in Fig. 1A) to the 3’RR creates a chromatin loop that mediates the subsequent synapsis of donor and acceptor S regions. In this manner, transcriptional activation and S-S synapsis are topologically coupled within the context of a single multiway 3D conformation (*12, 13, 25*) (Fig. 1A, right panel). In particular, it was proposed that within this multiway conformation (called a CSR center, CSRC), cohesin is loaded at the activated acceptor I promoter (I*γ*1 in Fig. 1A) leading to the formation of a new sub-loop via one-sided loop extrusion which leads to synapsis by aligning donor and acceptor S regions in the proximity of the 3’RR (Fig. 1A, right panel) (*12, 13*).

This mechanistic model of CSR, however, remains speculative. First, higher-order, multiway interactions at *Igh* have not been directly demonstrated, but rather, have been inferred from various chromosome conformation capture assays which mostly (4C-seq, Hi-C) or exclusively (3C) report pairwise interactions (*12, 25–27*). Therefore, it remains unknown whether the proposed multiway topologies exist during CSR and, if so, what their composition is. Second, recent work shows that cohesin does not load at promoters or enhancers, as was previously believed, but can be impeded near the 5’ and 3’ ends of active genes (*2*). Thus, the mechanism by which loop extrusion may drive synapsis and the precise contribution of S region transcription in this process requires further investigation.

In this study, we employ high-resolution Tri-C in CSR-activated primary murine B cells to unravel the multiway interactome on single *Igh* alleles from different viewpoints. We find that the 3’RR forms a self-interacting domain that, in contrast to the prevailing model, only infrequently associates with synapsed S regions and does not serve as a major loop anchor. Rather, S-S synapsis is most frequently aligned with the 3’CBE arguing that transcriptional activation by the 3’RR and S-S synapsis occur through different 3D conformations, that is, they are topologically uncoupled. Furthermore, we provide evidence that the conflict between loop extrusion and the high density of transcriptional complexes at activated acceptor S regions creates a CTCF-independent topologically associating domain (TAD) boundary spanning the entire acceptor S region. Consequently, S-S synapsis occurs at the interface of, and is underpinned by, this TAD boundary. Thus, our study reveals how structural changes in chromatin arising from conflicts between transcription and loop extrusion impact directly on the mechanism of a major biological process like CSR.

## RESULTS

### S-S synapsis is observed predominantly in proximity of the 3’CBE and is topologically uncoupled from 3’RR-mediated transcriptional activation

The first goal of this study was to determine whether multiway interactions proposed from pairwise interaction studies could be detected at the murine *Igh* locus during CSR. Hence, we used Tri-C, a chromosome conformation capture method that reports multiway interactions from a chosen viewpoint on single alleles (*29*) (Fig. S1A). In particular, Tri-C has successfully identified regulatory higher-order 3D hubs formed via the association of multiple enhancers and promoters (*29, 30*) which is not possible with 3C or 4C-seq approaches that detect mostly pairwise interactions. We performed Tri-C in primary, resting AID*^−/−^* B cells as well as primary AID*^−/−^* B cells activated with lipopolysaccharide (LPS), Interleukin 4 (IL4) and RP105 for 48h or 72h. This treatment primarily induces germline transcription of *Ighg1* from the I*γ*1 promoter and, consequently, IgG1 CSR (Fig. 1A). Since AID plays no role in enhancer-promoter interactions (*28, 31*), the use of AID*^−/−^* cells allowed us to evaluate locus topology in the absence of allelic rearrangements resulting from CSR. We found that although both 48h and 72h activation conditions yielded similar results, as seen by the high Pearson correlation coefficients between these datasets (Fig. S1B), enrichment of focal contacts was greater at 72h, which permitted better visualization of the data. Hence, we discuss the results from resting and 72h activated B cells below.

Since CSR in LPS + IL4 stimulated cells has been proposed to occur via multiway interactions between E*μ*, S*μ*, S*γ*1 and the 3’RR, we designed the following Tri-C probes: (1) A probe at the E*μ* enhancer (E*μ* capture fragment, Fig. 1B-C). S regions are repetitive and, therefore, it is not possible to design a probe within them. Since E*μ* is ~700 bp from S*μ*, we use the measurements from the E*μ* probe to infer contacts with S*μ* (*25–27*). (2) A probe at the hs4 enhancer of the 3’RR (hs4 capture fragment, Fig. 1D-E). (3) A set of four contiguous probes located <1kb from each other and spanning the I*γ*1 promoter (I*γ*1 capture fragment, Fig. 2A-B). This was done to increase the resolution of the assay from this viewpoint. The most distal I*γ*1 capture probe is ~1 kb from S*γ*1, hence we use the I*γ*1 capture measurements to infer contacts with S*γ*1 upon activation.

**Figure 2:**
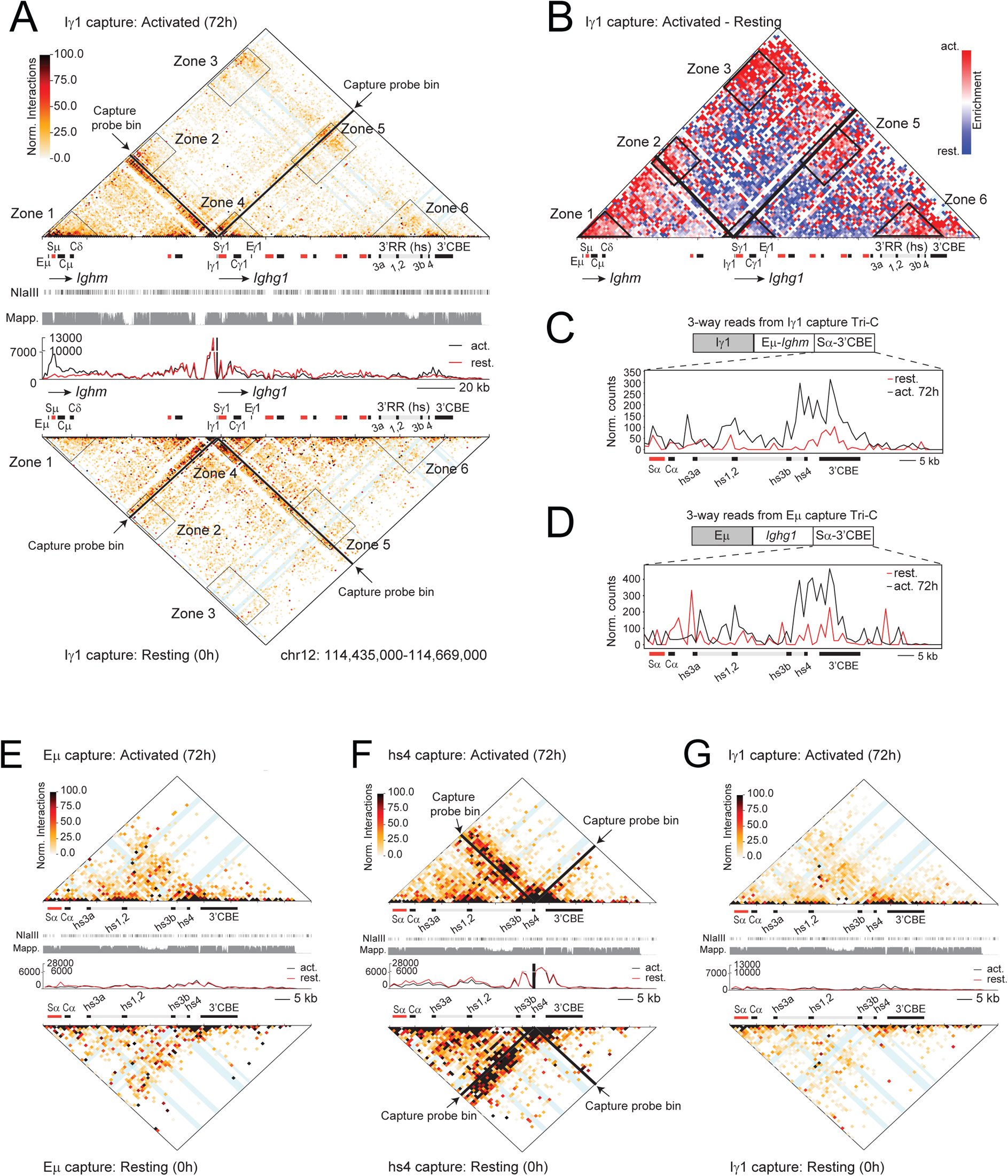
Multi-way interactions of the Iγ1 promoter in resting and CSR-activated B cells. A. Tri-C analysis from the Iγ1 promoter viewpoint in resting and activated B cells. The Iγ1-containing bins are blanked. Zones 1-5 are the same as in Fig. 1B with zone 3 (Iγ1-*Ighm*-3’RR-CBE) and zone 6 (Iγ1-3’RR-3’CBE) representing 3-way interactions.
B. Difference matrix (activated - resting) for the data shown in A. Colored bins show enriched contacts in activated cells (red) or resting cells (blue).
C. Histograms showing the distribution of contacts in the Sα-3’RR-3’CBE region obtained from three-way reads from the Iγ1 viewpoint (diagram above the bar plot). All reads contain the Iγ1 capture fragment, one reporter fragment mapping in the Eμ-*Ighm* region and the second reporter fragment aligning to the Sα-3’RR-3’CBE region. The cumulative frequencies of Sα-3’RR-3’CBE reporter reads are displayed in the histogram.
D. Histograms as in D above showing the distribution of contacts in the Sα-3’RR-3’CBE region obtained from three-way reads from the Eμ viewpoint (diagram above the bar plot). All reads contain the Eμ capture fragment, one reporter fragment mapping to the *Ighg1* region and the second reporter fragment aligning to the 3’RR-3’CBE region. The cumulative frequencies of Sα-3’RR-3’CBE reporter reads are displayed in the histogram.
E. Zoomed in view of the 3’RR-3’CBE region in resting and activated B cells from the Eμ capture Tri-C matrix in Fig. 1B.
F. Zoomed in view of the 3’RR-3’CBE region in resting and activated B cells from the hs4 capture Tri-C matrix in Fig. 1D.
G. Zoomed in view of the 3’RR-3’CBE region in resting and activated B cells from the Iγ1 capture Tri-C matrix in Fig. 2A.

Multiway reads in Tri-C libraries are defined as those that contain the capture fragment linked to 2 (3-way read) or more than 2 (>3-way read) additional fragments, termed reporter fragments (Fig. S1C). The reporter fragments harbor information on the genomic regions associated with the capture viewpoint. Therefore, all reporter fragments part of ≥3-way reads were extracted and displayed on a matrix. Since all reads contain the capture fragment, the bin harboring this viewpoint fragment is excluded in all matrices (indicated as a black stripe) (Fig. 1B, 1D and 2A). All matrices are normalized to contain 300,000 total reads which allows us to compare relative enrichments across the locus between different conditions (Methods). Since each multiway interaction is derived from single alleles, it provides information on the relative frequencies of the different 3D topological states of the locus within the population (*29*). It is important to note that the matrices in this study consist of 1 kb bins which allows us to delineate interactions between and within *Igh* genes (*Ighm,* 8.5 kb and *Ighg1,* 12.4 kb) and regulatory elements (3’RR, 27 kb and 3’CBE, 9.4 kb) at much greater resolution than previous studies employing 10 kb or larger bin sizes (*15, 26, 27, 32*).

We first asked if the known pairwise interactions between E*μ*, *Ighg1* and 3’RR, reported previously (*12, 25–27*), were recapitulated by Tri-C. In resting (0h) cells, E*μ* engages with the neighboring *Ighm* locale (zone 1) and also with the 3’RR-3’CBE segment (zone 3) forming a basal loop (Fig. 1B). Upon activation (72h), E*μ* and *Ighm* associate with transcribed *Ighg1* (zones 2 and 4) making contacts that spread across *Ighg1* (Fig. 1B-C). Similarly, from the 3’RR hs4 enhancer viewpoint, the basal loop between the 3’RR and E*μ* (zone 3) and the activation-induced 3’RR-*Ighg1* interactions (zones 4 and 5) were observed (Fig. 1D-E). Finally, using the I*γ*1 promoter as the viewpoint, we observed activation-dependent contacts of the I*γ*1 locale with *Ighm* (zones 1 and 2) indicative of S-S synapsis, and with the 3’RR-3’CBE segment (zones 5 and 6) (Fig. 2A-B). We conclude that Tri-C recapitulates the known pairwise interactions previously reported in CSR-activated B cells.

We next asked the key question that motivated the Tri-C analysis, namely, whether multiway interactions involving S-S synapsis with the 3’RR, as suggested from pairwise interaction studies, were observed in activated B cells. Indeed, zone 5 of the E*μ* capture matrix reveals a 3-way interaction between the E*μ* locale, *Ighg1* and the 3’RR-3’CBE segment in activated cells (Fig. 1B-C). Zone 2 of the hs4 capture matrix also reveals a 3-way interaction between the hs4, *Ighg1* and *Ighm* in activated cells (Fig. 1D-E). Surprisingly, however, closer inspection of the E*μ* capture matrix shows that zone 5 predominantly consists of contacts between the E*μ* locale, the entire *Ighg1* and the hs3b-3’CBE segment, with much weaker contacts involving the hs3a and hs1,2 enhancers and still weaker contacts in the intervening quasi-palindromic sequences between the hs3a and hs3b enhancers (Fig. 1B, zone 5). Since the hs3a-hs3b region spans ~22 kb whereas the hs3b-hs4 segment spans only ~5 kb, these results suggest that most of the 3’RR infrequently aligns to synapsed S regions. In fact, inspection of zone 5 from all capture viewpoints revealed that multiway interactions of *Ighg1* are strongly biased towards the hs3b-3’CBE segment relative to the hs3a-hs3b segment (Fig. 1B, D and Fig. 2A). In support, I*γ*1 capture analysis revealed that 3-way interactions involving the I*γ*1 locale and *Ighm* were more frequent with the hs3b-3’CBE segment than with the hs3a-hs3b segment (zone 3, Fig. 2A-B). We conclude that 3-way interactions involving synapsed S regions occur during CSR but that, in contrast to previous models (*12, 25*), S-S synapsis occurs mostly in proximity to the 3’CBE rather than the 3’RR.

To accurately determine the distribution of 3’RR-3’CBE contacts with synapsed S regions, we extracted from the I*γ*1 capture matrix all 3-way reads containing the I*γ*1 capture fragment, one reporter fragment mapping within the E*μ*-*Ighm* segment (spanning ~20 kb) and a second reporter fragment mapping to the S*α*-3’RR-3’CBE segment (Fig. 2C, upper panel). Similarly, from the E*μ* capture matrix, we extracted 3-way reads containing the E*μ* capture fragment, one reporter fragment mapping to the *Ighg1* segment and another reporter fragment mapping to the S*α*-3’RR-3’CBE region (Fig. 2D, upper panel). From these reads, the reporter fragments aligning to the S*α*-3’RR-3’CBE region were plotted to obtain a histogram revealing the distribution of reads in this region (Fig. 2C-D, lower panels). The interaction profile clearly revealed stronger contact frequencies in the hs3b-3’CBE segment compared to the hs1,2 and hs3a enhancers (Fig. 2C-D). Thus, within the CSRC, multiway S*μ*-S*γ*1-3’CBE contacts are more frequent than S*μ*-S*γ*1-3’RR contacts. These results further strengthen the conclusion that S-S synapsis is mostly associated with the 3’CBE rather than the 3’RR, as was formerly proposed.

The surprising under-representation of 3’RR interactions with the S-S synapsis state led us to inspect the 3’RR locale in more detail. Strikingly, Tri-C matrices showed that from all capture viewpoints, the 3’RR forms a self-interacting topological domain which associates with E*μ* or I*γ*1 capture viewpoints as an independent zone of multiway interactions (zone 6, Fig 1B and Fig. 2A, and magnified views in Fig. 2E-G). Importantly, interactions within the 3’RR locale include contacts with the intervening sequences between the hs enhancers, implying that the lower density of contacts in the hs3a-hs3b segment in zone 5 (Fig. 1B, D and Fig. 2A) is not due to poor mappability (Fig. 2E-G). Moreover, contacts between the individual 3’RR enhancers, especially between hs1,2 and hs3b-hs4, are discernible, suggesting the presence of internal secondary structures within the 3’RR that may bring the individual enhancers into proximity for synergistic action (Fig. 2E-G). We conclude that the 3’RR exists predominantly as a stable, self-associating domain at steady state.

Altogether, the Tri-C analyses show that the 3’RR self-interacting domain associates with the *Igh* promoters as a distinct topological state (zone 6 in Fig. 1B, D and Fig. 2A) and that S-S synapsis occurs as a separate state associated mostly with the 3’CBE (zone 5 in Fig. 1B and zone 3 in Fig. 2A). We infer that these represent two distinct topologies of the locus within the population that, mechanistically, reflect the uncoupling of 3’RR-mediated transcriptional activation from 3’CBE-anchored S-S synapsis.

### Transcription at *Ighg1* creates a CTCF-independent *de novo* TAD boundary that underpins S-S synapsis

Upon closer inspection of the Tri-C cumulative contact frequency profiles, we observed an asymmetry of contact distribution at *Ighg1* (Fig. 3A). From the E*μ* viewpoint, contacts were higher upstream of C*γ*1 with a substantial decay within and downstream of C*γ*1 (Fig. 3A, arrow). From the hs4 viewpoint, however, a large spread of contacts is observed within and downstream of C*γ*1 with substantially fewer interactions upstream of C*γ*1 (Fig. 3A). These distinct viewpoint-specific asymmetric signatures indicate that *Ighm* is more engaged with sequences upstream than downstream of C*γ*1. In contrast, the 3’RR-3’CBE region is more engaged with sequences downstream than upstream C*γ*1 (Fig. 3A). Additionally, the I*γ*1 viewpoint, located upstream of S*γ*1, contacted the E*μ*-C*μ* region more frequently than the proximal C*γ*1 region (Fig. 3A). These observations led us to hypothesize that in activated B cells, *Ighg1* acts as a topological barrier for chromatin interactions, akin to a TAD boundary.

**Figure 3:**
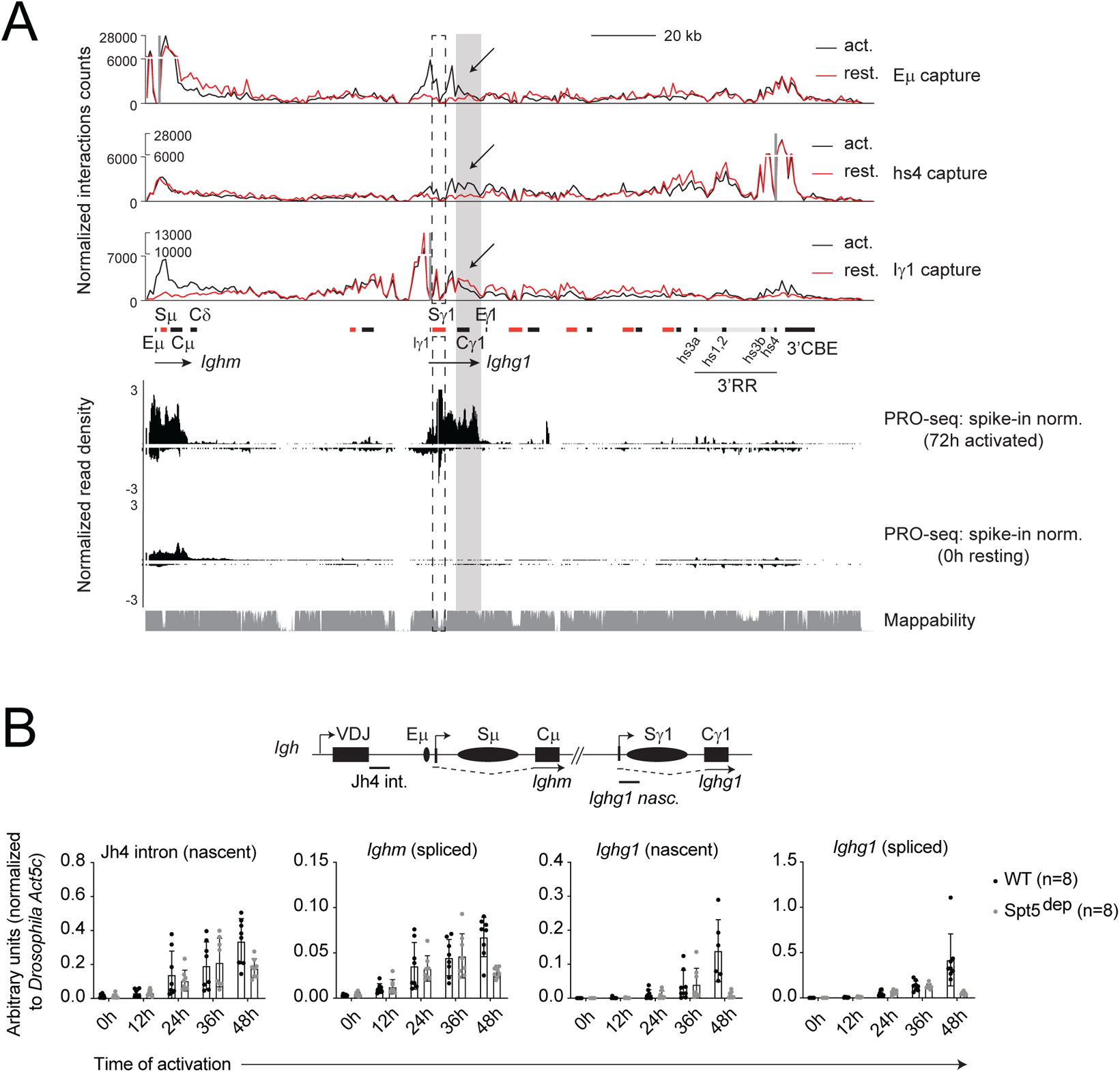
Asymmetric contact frequencies at *Ighg1*: evidence for a topological boundary in activated cells. A. Cumulative frequency histograms of Tri-C data from Eμ (Fig. 1B), hs4 (Fig. 1D) and Iγ1 (Fig. 2A) viewpoints assembled together to show the symmetry of contact distribution at *Ighg1*. The highly repetitive Sγ1 showing reduced mappability is highlighted with a dashed box. Cγ1 and a downstream portion is shaded in grey and the arrows highlight the viewpoint-specific asymmetry of contact frequencies upstream and downstream of Cγ1, as described in the text. The nascent transcription data obtained by PRO-seq in resting and activated cells were normalized to reads from spiked-in *Drosophila* S2 cells before alignment to the mouse genome.
B. RT-qPCR analysis of nascent and spliced *Igh* transcripts in WT and Spt5^dep^ cells at the indicated time points post-activation. B cells were mixed with *Drosophila* S2 cells prior to lysis and RNA extraction. The qPCR data was normalized to the levels of the *Drosophila* housekeeping gene, *Act5c*. The location of the transcripts analyzed is shown in the diagram above the graphs.

TAD boundaries are frequently formed at DNA elements bound by the protein CTCF, which acts as a barrier for cohesin-containing loop extrusion complexes (*8, 32, 33*). However, with the exception of C*α*, which harbors a single CTCF site, there are no CTCF sites in the *Igh* genes(*34*), which rules out CTCF as the molecular basis for TAD boundary formation at *Ighg1*. The key difference between resting and activated B cells is the presence of robust transcription at *Ighg1*, as seen by precision run-on sequencing (PRO-seq) analysis (Fig. 3A). Moreover, B cell activation is associated with global transcriptional amplification(*35*) that also results in a major upregulation of transcription at *Ighm* which, importantly, is only evident when the data is normalized to reads from externally spiked-in *Drosophila* S2 nuclei (Fig. 3B) but not when normalized internally to the *Gapdh* transcript (Fig. S1D). Thus, we asked if transcription at *Ighg1* could, by itself, create a new TAD boundary in activated cells leading to the formation of *Igh* sub-TADs.

To test this idea directly, we performed Tri-C from an intergenic viewpoint downstream of *Ighg2a* (henceforth called *Ighg2a*), a transcriptionally silent region which does not engage in focal contacts with E*μ*, I*γ*1 or hs4 viewpoints (Fig. 1B, D and Fig. 2A). We reasoned that if the transcribed S*γ*1 region acts as a TAD boundary, then *Ighg2a* would interact more frequently with sequences upstream of *Ighg1* in resting cells (where S*γ*1 in silent) than in activated cells. Indeed, Tri-C revealed that in resting cells, *Ighg2a* contacts spanned the entire locus, whereas in activated cells, contacts were largely restricted within the *Ighg1*-3’CBE domain harboring the capture fragment (Fig. 4A). We note that the sparse appearance of the matrix in activated cells is not due to lower sequencing depth (Fig. S1C), and moreover, all matrices contain 300,000 normalized reporter reads. As seen in the cumulative contact frequency histogram, the loss of distal contacts in activated cells are compensated by a gain of interactions in the viewpoint locale (Fig. 4A).

**Figure 4:**
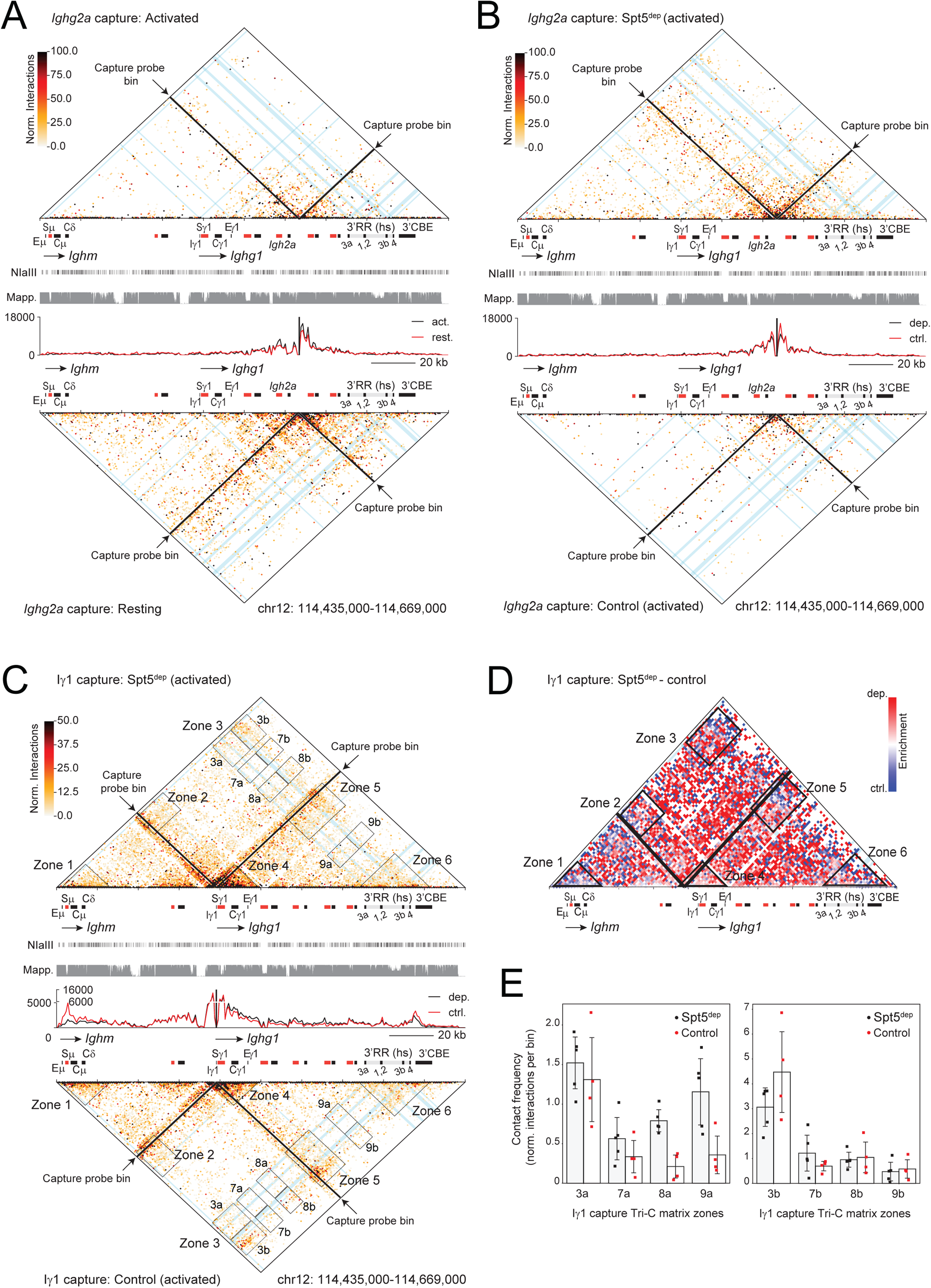
Transcription at *Ighg1* creates a *de novo* TAD boundary in activated B cells. A. Tri-C analysis from a transcriptionally inert intergenic region downstream of *Ighg2a* in resting and activated B cells.
B. Tri-C with the *Ighg2a* viewpoint as in A but from control and Spt5^dep^ activated B cells.
C. Tri-C analysis from the Iγ1 promoter viewpoint in *Rosa26*^Cre-ERT2/+^ (control) and *Supt5h*^F/−^*Rosa26*^Cre-ERT2/+^ (Spt5^dep^) activated B cells. The boxed zones 1-6 are the same as in Fig. 2A. Additionally, zone 3 is split into zones 3a and 3b to distinguish between hs3a-hs3b (3a) and hs3b-3’CBE (3b) segments. Moreover, three zones representing non-specific multiway contacts between the Iγ1 viewpoint and the 3’RR-3’CBE are indicated (zones 7, 8, 9). These are further divided based on interactions involving the hs3a-hs3b segment (zones 7a, 8a, 9a) or the hs3b-3’CBE segment (zones 7b, 8b and 9b).
D. Difference matrix (Spt5^dep^ - control) for the data show in C.
E. Bar plots quantifying the interaction frequency (contacts per bin) within the zones indicated in the Iγ1 capture matrices in C. Each dot represents replicate samples from different experiments. The left plot shows interactions in zones 3a, 7a, 8a and 9a and the right plot shows contacts in zones 5b, 7b, 8b and 9b.

Next, we hypothesized that if a high density of transcriptional complexes at S*γ*1 could serve as a physical impediment to loop extrusion complexes, then loss of *Ighg1* expression in activated B cells should alleviate this blockade. To address this, we took advantage of our previously reported system where depletion of the transcriptional elongation factor, Spt5 (Spt5^dep^), causes a ~10-fold decrease in *Ighg1* transcription (Fig. 3B) along with decreased long-range interactions of *Ighg1* in activated B cells measured by 3C-qPCR (*36*). In this system, the floxed *Supt5h* allele, encoding Spt5, is combined with the Cre recombinase expressed constitutively from the *Rosa26* promoter (*Supt5h*^F/−^ *Rosa26*^Cre-ERT2/+^, Spt5^dep^) (*37*). Upon addition of 4-hydroxytamoxifen (4-HT), excision of the floxed segment leads to depletion of Spt5 protein. As a control, we use *Rosa26*^Cre-ERT2/+^ cells treated with 4-HT (henceforth termed control). Of note, Spt5 depletion in this system does not have a major impact on global gene expression (*36*). In essence, this system allowed us to ask whether loss of *Ighg1* transcription under activating conditions abolished topological insulation at *Ighg1*.

The inert *Ighg2a* viewpoint made contacts mostly within the *Ighg1*-3’CBE TAD in control cells (similar to the activated cells in Fig. 4A). Strikingly, in Spt5^dep^ cells, contacts extended well beyond *Ighg1* (Fig. 4B). The cumulative contact frequency histogram reveals how interactions in the *Ighg2a* capture viewpoint locale are higher in control cells than in Spt5^dep^ cells (Fig. 4B). Indeed, the *Ighg2a* contact matrix in activated Spt5^dep^ cells is virtually identical to the *Ighg2a* matrix in resting B cells where *Ighg1* is also silent (compare Fig. 4A lower matrix with Fig. 4B upper matrix). These findings are consistent with the loss of a TAD boundary in activated Spt5^dep^ cells. We note that creating a sub-TAD can interfere with gene expression within the larger, original TAD by blocking enhancer activity (*38–42*). However, the robust upregulation of *Ighm* transcription upon activation (Fig. 3A-B) implies that the TAD boundary at *Ighg1* does not act as a topological barrier for the 3’RR to regulate the I*μ* promoter.

We next performed Tri-C from the I*γ*1 viewpoint, located upstream of the putative TAD boundary in S*γ*1, in control and Spt5^dep^ cells to determine whether the multiway conformation mediating S-S synapsis was affected by the loss of the TAD boundary at *Ighg1*. Indeed, we observed decreased multiway contacts between the I*γ*1 locale, *Ighm* and the 3’CBE in Spt5^dep^ cells (zone 3, Fig. 4C-D) as well as between the I*γ*1 locale and *Ighm* (zone 1, Fig. 4C-D). In addition, contact densities were visibly increased downstream of the I*γ*1 capture viewpoint which is also evident from the cumulative contact frequency histogram (Fig. 4C). Indeed, analysis of I*γ*1 capture Tri-C in Fig. 2A shows that resting B cells (which lack *Ighg1* transcription) harbor increased non-specific contacts downstream of I*γ*1 compared to activated cells (Fig. 2A matrix and cumulative frequency histogram).

To quantify non-specific multiway contacts with the I*γ*1 locale, we compared the changes in contact density in Spt5^dep^ cells in zone 3 as well as three locations on the matrix corresponding to multiway interactions between the I*γ*1 viewpoint, the 3’RR-3’CBE segment and random locations (zone 7, 8, 9; Fig. 4C). Since, as discussed above, most of the 3’RR is not part of multiway interactions with synapsed S regions, we further divided the 3’RR-3’CBE region into two segments: (a) from hs3a to the 5’ end of hs3b (zones 3a, 7a, 8a, 9a), and (b) from hs3b to the end of the 3’CBE (zones 3b, 7b, 8b, 9b) (Fig. 4C). The resulting bar graphs show that contacts are reduced in the hs3b-3’CBE segment (zone 3b) in Spt5^dep^ cells, corresponding to the decrease in multiway contacts underlying S-S synapsis, but contacts are largely unaffected in zones 7b, 8b and 9b (Fig. 4E). In contrast, multiway contacts of I*γ*1 with the hs3a-hs3b segment show increased contacts with all chosen locations in Spt5^dep^ cells with the strongest increases seen in the non-specific zones (7a, 8a, 9a) (Fig. 4E). Thus, the loss of *Ighg1* transcription in Spt5^dep^ cells results in increased non-specific contacts across the locus. In fact, this can also be observed from the general increase (red bins) across *Igh* in the difference matrix (Fig. 4D).

Collectively, these results provide further support for the notion that transcriptional activity creates a *de novo* TAD boundary at *Ighg1* in activated B cells that impedes loop extrusion across *Ighg1.* S regions are enriched in slowly elongating and paused RNA polymerase II (*43–45*). Hence, we infer that the high density of transcription complexes creates a potent and stable loop extrusion barrier that spans the entire length of the S region. As a result, S-S synapsis occurs at the interface of, and is dependent upon, the creation of this TAD boundary.

### Micro-C analysis provides further evidence for a transcription-dependent TAD boundary at *Ighg1* underpinning S-S synapsis

Since Tri-C provides locus-specific views of contact frequencies, we asked whether similar results would be observed via an alternative approach that captures all genomic interactions without locus enrichment. Hence, we performed Micro-C, which reports genome-wide interactions between any pair of loci, akin to Hi-C, but at much higher resolution due to its use of micrococcal nuclease to digest crosslinked chromatin (*46*). Micro-C contact matrices were plotted in 1 kb bins (the same resolution as the Tri-C matrices) which allow for much finer resolution of contacts between *Igh* genes and regulatory elements than with previous Hi-C analyses.

In control cells, *Ighm* and *Ighg1* mostly interacted with the hs4-3’CBE region (boxes 1b and 2b in Fig. 5A) and form weaker interactions with the 3’RR (boxes 1a and 2a in Fig. 5A). Moreover, hs4-3’CBE contacts were spread across *Ighm* and *Ighg1* with no discernible bias at E*μ* or the I*γ*1 promoter (Fig. 4A, boxes 1b and 2b). Strikingly, the 3’RR forms a distinct, self-interacting domain that appears to be bounded by a loop between C*α* and the 3’CBE (dashed circle labeled as 4 in Fig. 5A). It is plausible that this loop is formed between the known CTCF-bound site in C*α* (*47*) and the CTCF sites in 3’CBE pointing in convergent orientations as seen at many TAD boundaries genome-wide(*32*). Importantly, the substantial enrichment of contacts within the 3’RR is in contrast with the relatively infrequent contacts of the 3’RR with *Ighm* and *Ighg1* (boxes 1a and 2a). This observation resembles our Tri-C results (Fig. 1B, D and Fig. 2A) in showing that the 3’RR exists as a topologically distinct unit that, relative to the 3’CBE, engages in infrequent steady-state contacts with *Ighm* and *Ighg1.* Finally, and most strikingly, interactions of *Ighm* with *Ighg1* were strongest in the I*γ*1-S*γ*1 region whereas C*γ*1 was relatively depleted of contacts with *Ighm* (Fig. 5A, box 3). Conversely, interactions of the hs4-3’CBE region were strongest in C*γ*1 and downstream sequences and relatively weaker in the I*γ*1-S*γ*1 region (Fig. 5A, box 2b). These results reveal a clear asymmetry of contacts at *Ighg1* as seen also in Tri-C (Fig. 3A) and support the conclusions drawn from Tri-C that *Ighg1* serves as a bona fide TAD boundary at *Igh* in activated B cells.

**Figure 5:**
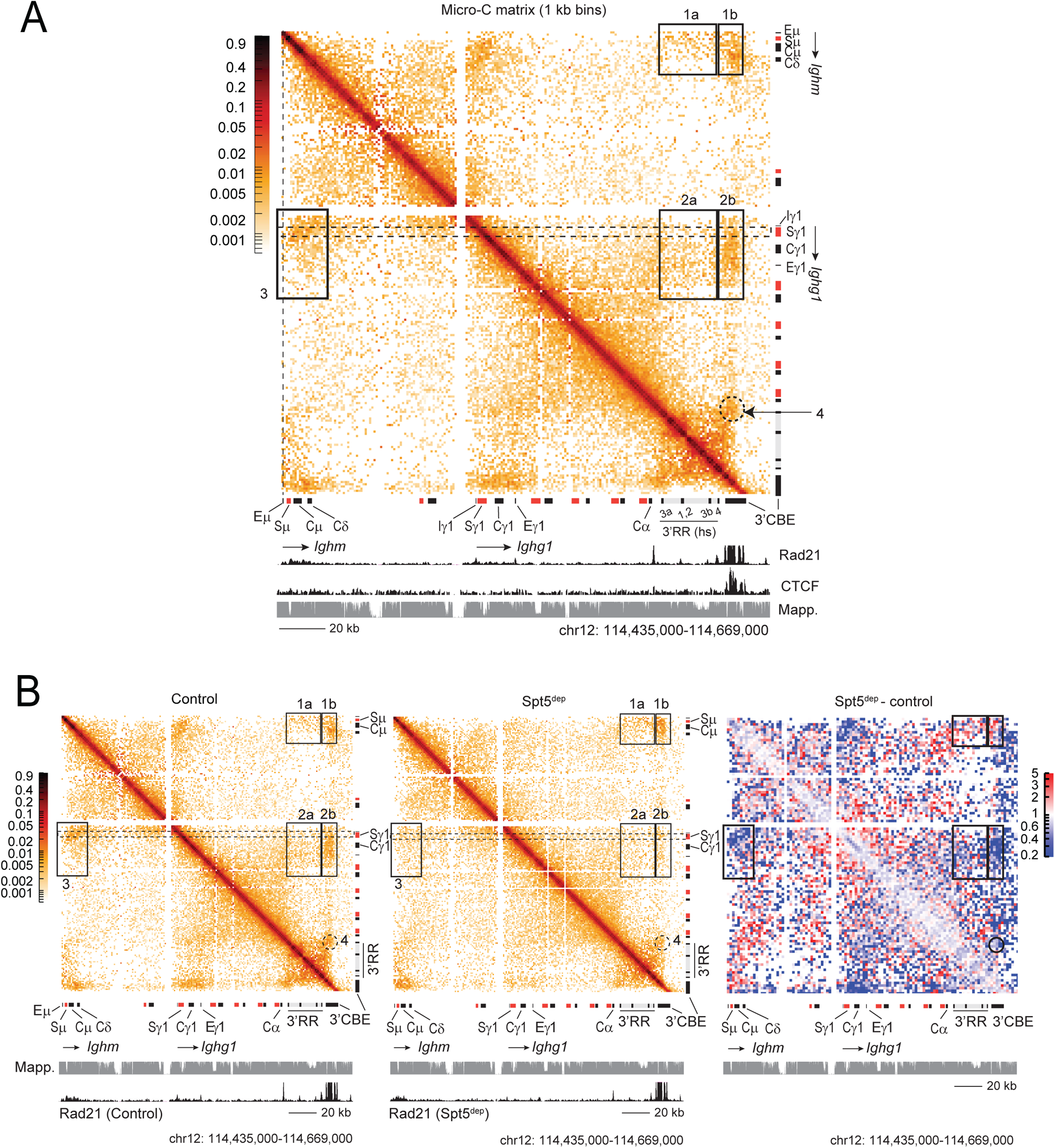
Micro-C reveals a transcription-dependent TAD boundary at *Ighg1* and a CTCF-bounded self-interacting 3’RR domain. A. Micro-C matrix in activated *Rosa26*^Cre-ERT2/+^ (control) B cells at the *Igh* locus resolved in 1 kb bins, the same resolution as the Tri-C matrices. Key insights gained from Micro-C, particularly the similarities with Tri-C, are highlighted and described in the main text. Boxed regions correspond to *Ighm*-3’RR-3‘CBE contacts (boxes 1a, 1b), *Ighg1*-3’RR-3’CBE contacts (boxes 2a, 2b), *Ighg1*-*Ighm* contacts (box 3). The Cα-3’CBE loop is encircled (circle 4). Rad21 ChIP-seq track represents cohesin occupancy. The CTCF track in activated B cells is from Thomas-Claudepierre et al. (Ref. 27). The mappability track is in grey. The white stripes on the matrix correspond to poorly mappable sequences.
B. B. Micro-C analysis from activated *Rosa26*^Cre-ERT2/+^ (control) (left matrix, identical to the matrix in A above) and activated *Supt5h*^F/−^*Rosa26*^Cre-ERT2/+^ (Spt5^dep^) (middle) cells. Boxed and encircled regions and tracks below the matrices are as described in A. The difference matrix (Spt5^dep^ - control), binned at 2 kb resolution, is shown on the right where red and blue bins indicate enrichments in Spt5^dep^ cells and control cells, respectively.

Micro-C analysis from Spt5^dep^ cells revealed that *Ighg1* interactions with *Ighm* (that is, S-S synapsis) and the hs4-3’CBE region were diminished in these cells (blue bins in Fig. 5B). This was accompanied by an increase in long-range interactions across the *Igh* locus (red bins away from the diagonal in Fig. 5B, right) as well as within the *Ighg1* locale (red bins close to the diagonal at *Ighg1* in Fig. 5B, right). The results are consistent with the loss of a TAD boundary at *Ighg1* in Spt5^dep^ cells and validate the results and conclusions drawn from Tri-C. Of note, occupancy of Rad21, a cohesin subunit, at the 3’CBE and *Ighm* is unchanged in Spt5^dep^ cells but is reduced in *Ighg1* concomitant with decreased transcription (Fig. 5B). Moreover, Rad21 is reduced at C*α* in Spt5^dep^ cells resulting in a slight loss of insulation of the 3’RR self-interacting domain (Fig. 5B, circle 4).

We conclude that transcription at *Ighg1* functions as an efficient and robust TAD boundary in activated B cells that underpins the S-S synapsis state critical for CSR.

## Discussion

Our study harnesses the power of Tri-C to identify higher order interactions on single alleles to unravel key principles of chromosome conformation during CSR. Firstly, we report that multiway interactions between synapsed S regions and regulatory elements can be detected at steady state in activated B cells. Secondly, the high-resolution of Tri-C allows us to determine the composition of the different 3D conformations at steady-state revealing new and unexpected insights into the mechanism of CSR. Finally, we identify a transcription-dependent TAD boundary in activated B cells at acceptor S regions that underpins the S-S synapsis conformation critical for CSR. Although it is established that transcription can majorly influence chromatin structure around active genes and create topological barriers by modulation of the loop extrusion machinery (*1–6*), whether such conflicts between loop extrusion and transcription contribute directly to biological processes has not been described. Thus, from a conceptual viewpoint, our study highlights the fact that the *de novo* creation of a TAD boundary by transcriptional activity directly regulates the mechanism of a major physiological process, namely, CSR.

Our results lead to a revised model of CSR (Fig. 6). Naïve, resting B cells harbor a dynamic, basal loop between the 3’RR-3’CBE and the entire E*μ*-*Ighm* regions with no specific bias for E*μ* or 3’RR as barriers to cohesin complexes, in line with our new results (step 1, Fig. 6). Upon B cell activation, loop extrusion brings the 3’RR and I*γ*1 promoter, the latter primed by transcription factors, into proximity to activate *Ighg1* transcription (step 2, Fig. 6). B cell activation also upregulates transcription at *Ighm* as part of the global amplification of the B cell transcriptome (*35*). Transcription results in a high density of Pol II complexes particularly in S*γ*1 and S*μ* where secondary structures cause Pol II pausing (*43–45*) (step 3, Fig. 6). Once high rates of transcription are established, a TAD boundary is created at S regions. This boundary acts as a barrier to subsequent loop extrusion resulting in the synapsis of S*μ* and S*γ*1 (step 4). Subsequent steps such as AID targeting, DNA break formation and ligation lead to deletional CSR (Fig. 6).

**Figure 6:**
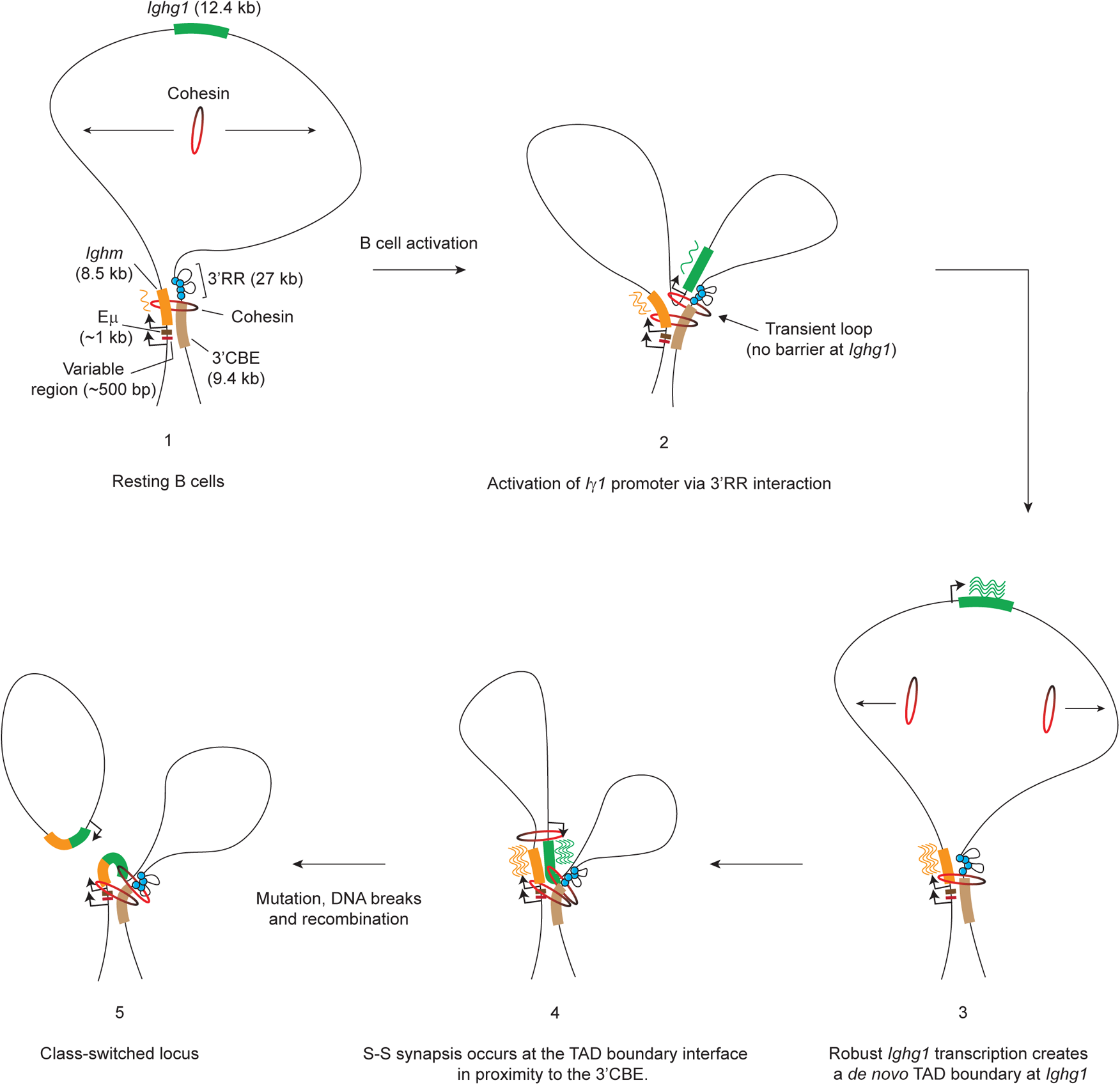
Mechanistic model for the role of transcription-dependent TAD boundary creation in S-S synapsis. Model for IgG1 CSR incorporating insights from Tri-C and Micro-C and described in detail in the Discussion. The 3’RR is shown as a self-interacting domain which brings the hs sites (blue circles) in closer proximity to each other. A basal loop exists between the Eμ-*Ighm* region and the 3’CBE in resting B cells (step 1). Based on our findings here, we propose that this loop forms via a transcription-mediated loop extrusion barrier at *Ighm*, especially at Sμ where Pol II pausing is elevated. Our data suggest that Eμ only makes a minor contribution to this basal loop which suggests that Eμ is not a major anchor for the loop, consistent with Eμ being dispensable for robust CSR. Upon activation, cohesin loads at a random location and loop extrusion brings the 3’RR in proximity of *Ighg1* to activate the Iγ1 promoter (step 2). Since there is no transcription at *Ighg1* at this stage, there is no barrier to loop extrusion. Hence, such interactions are likely to be transient resulting in dissociation of this loop (step 3). Once high rates of transcription are achieved (step 3), a robust TAD boundary is created that impedes subsequent rounds of loop extrusion resulting in Sμ-Sγ1 synapsis at the TAD boundary interface. Importantly, we propose, based on our results here, that S-S synapsis is predominantly supported by multiway interactions with the 3’CBE and not the 3’RR as had been previously proposed (step 4). At this stage, AID-induced DNA breaks in S regions and their ligation via non-homologous end-joining leads to deletional recombination and productive CSR to express IgG1 (step 5). Importantly, although such a representation can only show static interactions, our data imply that all such interactions are dynamic. For example, the 3’CBE engages along the entire length of *Ighm* and *Ighg1* (resulting in the appearance of “stripes” in the matrices) and indicative of a dynamic interaction between these regions within the cell population.

The model differs from previous models (*12, 13, 25*) in three important aspects. Firstly, the 3’CBE, rather than the 3’RR, is dynamically aligned with synapsed S regions (step 4, Fig. 6). The 3’RR forms a self-interacting domain that infrequently contacts the synapsed S regions. Secondly, following activation of the I*γ*1 promoter upon contact with the 3’RR (step 2, Fig. 6), transcription of *Ighg1* ensues during which time *Ighg1* is dissociated from the basal loop (step 3, Fig. 6). This is in contrast to the previous model wherein primed *Ighg1* is transcribed within the CSRC following entrapment with cohesin rings (*12, 13*). Our rationale is that since *Ighg1* is not transcribed at this stage, there is no topological barrier at *Ighg1* to impede loop extrusion and create a stable, multiway conformation. We reason that a certain period must lapse after enhancer-mediated activation during which *Ighg1* transcription reaches robust levels in order to effectively create a TAD boundary. Only then, via new loop extrusion events, can S-S synapsis occur at the interface of the TAD boundary (step 4, Fig. 6). This reasoning, together with the fact that we could not detect preferential enrichments of contacts between I*γ*1 (or I*μ*) with the 3’RR even at 1 kb resolution, suggests that the interaction of the 3’RR with the I*γ*1 promoter is transient. Therefore, we conclude that I promoter activation constitutes a separate topological state (step 2) from S-S synapsis (step 4) within the population (Fig. 6). This is borne out by our RT-qPCR results showing that maximal *Ighg1* transcription is reached 48h after B cell activation (Fig. 3B). One possibility is that transient contact of the 3’RR with the I*γ*1 promoter leads to transcriptional bursting which can result in high transcriptional activity in the absence of a stable enhancer-promoter loop.

The third important difference between ours and the previous model regards where cohesin loads on *Igh*. It was proposed that cohesin loads at *Igh* enhancers and the I*γ*1 promoter upon B cell activation (*12, 13*). This conclusion was based on ChIP data showing that the cohesin loader, Nipbl, was highly enriched at transcription start sites (*3, 12, 48–50*). However, as shown in a new study (*2*), the Nipbl antibody used in these studies was non-specific resulting in erroneous maps of Nipbl chromatin occupancy. Using a new epitope-and degron-tagging approach, Nipbl was detected only rarely at transcription start sites (*2*). With these new data and *in silico* modeling, the authors propose that cohesin loads randomly on chromatin and accumulates at gene termini and enhancers due to the physical barrier imposed by transcriptional complexes (*2*). Hence, we incorporate these crucial insights into our model by showing that cohesin loads randomly on *Igh* chromatin and is subsequently impeded by transcription at S regions to achieve S-S synapsis (Fig. 6).

E*μ* was suggested to act as a major cohesin loading site and loop extrusion barrier (*12, 13*). Yet, deletion of E*μ* results in a mild CSR phenotype (*51–53*). Close inspection of both Tri-C and Micro-C data reveals that contacts at *Ighm* broadly span the E*μ*-C*μ* region and extend into C*δ* with no discernible enrichment bias at E*μ*. This suggests that contacts within the S*μ*-C*μ* zone are sufficient to support S-S synapsis and 3’RR interactions, with E*μ* making only a minor contribution. This provides a plausible explanation for the weak CSR phenotype in E*μ*-deficient cells. In fact, *Ighm* harbors similar levels of nascent transcription (Fig. 3A) and cohesin (Rad21) (Fig. 5A) as *Ighg1*. This suggests that the basal loop between the *Ighm* region and the 3’CBE in both resting and activated cells may be formed when extruding cohesin complexes (loaded downstream of *Ighm*) are obstructed by transcriptional complexes at *Ighm,* particularly within the S*μ* repeats where Pol II pausing is elevated (*43–45*) (Fig. 6). Indeed, contact frequency histograms from the hs4 and I*γ*1 viewpoints show that interactions at *Ighm* peaked around S*μ* and decreased thereafter towards E*μ* (Fig. 3A). Thus, we propose that all chromatin loops involved in CSR are formed by conflicts between transcriptional and loop extrusion complexes.

Recently, it was reported that the deletion of all ten CTCF sites in the 3’CBE had, surprisingly, no impact on IgG1 CSR and variable effects on transcription and CSR of other isotypes (*54*). We suggest that the substantial contact density that we observe in the region just preceding the 3’CBE, that is, from 3’ of hs4 up to the 5’ boundary of the 3’CBE, is sufficient for anchoring long-range interactions during CSR in a CTCF-independent manner, perhaps mediated by transcription from hs3b and hs4. Finally, we note that the interaction between C*α* and the 3’CBE, most noticeable in the Micro-C analysis, has not been observed before, to our knowledge, but is in line with the fact that CTCF is bound to C*α* in resting, naive B cells and its occupancy is reduced, but not lost, upon activation (*12, 34*). Ablation of this site leads to premature activation of some *Igh* genes in resting B cells revealing its insulator function (*47*). Nevertheless, it is evident that an extruding cohesin loop would need to bypass CTCF bound to C*α* in order to generate the contacts between 3’CBE and *Igh* genes, suggesting that this CTCF site is permissive to extruding cohesin in activated B cells.

In conclusion, we expect that our work will impact on future studies not only in antibody diversification but in other genomic processes where loop extrusion and transcription play important roles, such as DNA repair. We propose that the use of high-resolution Tri-C and Micro-C provides a powerful combinatorial strategy to probe the finer details of 3D chromosome conformational changes underlying genomic interactions and thereby gain deeper insights into biological mechanisms.

## METHODS

### Mouse breeding and culturing of primary, splenic B cells

AID*^−/−^* mice (*20*) were maintained in a C57BL/6 background and housed in the IMBA-IMP animal facility in standard IVC cages with HEPA filtering. All experiments were conducted in compliance with IMP-IMBA animal facility guidelines, and Austrian and EU law. Primary B cells were prepared from spleens of 2–4-month-old mice and cultured in complete RPMI as per standard protocols (*31*). B cells were cultured in complete RPMI medium supplemented with 10% fetal bovine serum and antibiotics, Interleukin 4 (IL4; made in-house by the Molecular Biology Service, IMP), 25 *μ*g/ml Lipopolysaccharide (Sigma) and RP105 (made in-house). Transgenic *Supt5h*^F/−^*Rosa26*^Cre-ERT2/+^ (Spt5^dep^) and *Rosa26*^Cre-ERT2/+^ (control) mice were described previously (*36*). For these experiments, 30h after seeding, 2 *μ*M 4-Hydroxytamoxifen (Sigma) was added for 32h followed by harvesting.

### Tri-C

We performed TriC experiments as described (*30, 55*) with several modifications. We used 1.5 x 10^7^ resting, naïve splenic B cells (0h) and 10^7^ activated splenic B cells. Cells were washed twice in PBS containing 10% FBS (PBS/FBS) and crosslinked in 2% methanol-free Formaldehyde in PBS/FBS in 15 ml tubes at room temperature for 8.5 minutes with occasional, gentle inversion. The reaction was quenched with 0.133M Glycine followed by two washes in cold PBS/FBS (5 min, 1500 rpm for activated cells and 14,000 rpm for naïve cells). For cell lysis, the pellet was resuspended in 1ml PBS/FBS and 4ml C1 lysis buffer (0.32M Sucrose, 10mM Tris-HCl pH 7.5, 5mM MgCl_2_, 1% Triton X-100) with 1x protease inhibitors (Roche) and samples were rotated for 30 min at 4°C. Nuclei were centrifuged at 2000 rpm for 10 min at 4°C and the pellet was resuspended in 2 ml C1 lysis buffer with 1x protease inhibitors followed by another round of centrifugation. Nuclei were resuspended in 50µl 0.5% SDS and transferred to 1.5 ml DNA-low-bind safe-lock tubes and shaken at 500 rpm, 62°C for 5 min in a thermomixer. To sequester SDS, 145µl water and 25µl 10% Triton-X-100 were added to the tubes and incubated at 37°C for 15 min with shaking at 500 rpm. Next, 220µl water and 55µl 10x CutSmart Buffer (NEB) were added and 50-100µl of sample were transferred to a new tube (undigested control), the same volume was refilled with dH2O and 30 µl NlaIII (10U/µl, NEB) were added. The digest was incubated at 37°C, 500 rpm, shaking overnight. The next morning, an additional 30 µl NlaIII enzyme was added and incubation continued for 5-6h. Next, 50-100µl of the digested sample was transferred to a new tube (digested control) and the rest of the sample was heat-inactivated for 20 min at 65°C and briefly chilled on ice. Ligation with 10.000U of T4 DNA Ligase (NEB) in a total volume of 1275µl was performed at 16°C, 1400 rpm overnight.

Samples and controls were treated with 10µl and 1.5-3µl Proteinase K (20mg/ml, Sigma), respectively, and incubated at 65°C, 500rpm overnight. Next, 10µl and 2,5-5µl RNaseA (20mg/ml, Invitrogen), respectively, was added followed by incubation at 37°C for 30 minutes, 500 rpm. To prepare genomic DNA, the samples (split into two 2ml tubes) and controls were extracted once with phenol-chloroform-isoamyl alcohol, once with chloroform and then precipitated with 0.3M NaOAc and 100% Ethanol for 1h at −20°C. Samples were centrifuged at full speed at 4°C, pellets were washed twice in 70% Ethanol and dry pellets were resuspended in 100µl (samples) or 20µl (controls) dH2O. Sample integrity was checked on a 0.7% agarose gel, loading all of the controls and 5-10µl of the samples. DNA concentration was measured using the Qubit dsDNA HS assay kit (Invitrogen) and 2x 6µg of DNA in 105µl was used for sonication on Covaris S2 focused ultrasonicator (4°C, 10% duty, 5 intensity, 200 cycles/burst 50-55 sec, frequency sweeping) in Covaris microtubes (AFA-Fiber, snap cap). Following a 0.7x AmpureXP beads (Agencourt) purification, the sonicated material was checked on a fragment analyzer. Three samples of 2 µg each with a modal distribution of 450-500 bp were treated with NEBNext UltraII DNA Library Preparation kit for Illumina for end repair and adaptor ligation using 5µl of NEBNext Adaptor for Illumina (E7601A) and purified with 1.1x AmpureXP beads. To amplify all material, six PCR reactions, primers with Nextflex unique dual index barcode sequences (Perkin Elmer) were used and PCR reactions were purified with 1.1x AmpureXP beads followed by elution in 34µl dH2O. All samples were individually checked on a fragment analyzer for successful adapter addition and then pooled to obtain one library (with 6 barcode combinations) per original sample. Concentration was measured by Qubit. For capture enrichment with individual 5’biotinylated capture probes, 1-2µg DNA per library was pooled. All libraries for the same capture probe were first mixed and then split to a maximum of 6 libraries per tube. Capture hybridization was done using the HyperCap Target enrichment kit (Roche 08286345001) with 5µg of mouse COT DNA and 5µl universal blocking oligos per library. After 2x AmpureXP beads purification, samples were directly eluted from the beads into 7.5µl 2x hybridization buffer and 3µl hybridization component A from the kit per library. Next, 4.5µl of 2.9nM per capture probe per library was added. For the IgG1 pool, an 11.6nM capture probe mix of 4 probes was used. For hybridization, pull down, washing and PCR procedures, we followed the original protocol except for using standard Illumina P5 and P7 primers and 13 cycles for PCR. A 1.1x AmpureXP beads purification was performed and 33µl dH2O per captured library was used for elution. Using the pool of eluted libraries, a second round of hybridization was set-up, performed as above followed by PCR with 9-10 cycles using P5 and P7 primers. A 0.7x AmpureXP beads purification was applied and samples were eluted in 5µl water per library and concentrations of the pools were measured by Qubit. Finally, the purified pool of captured libraries was analyzed on a fragment analyzer. For multiplexed NGS analysis, samples of different captures were pooled in equimolar amounts and subjected to paired-end sequencing using at least 300 cycles on Illumina NextSeq or NovaSeq platforms.

### Micro-C

Micro-C was performed with 7×10^6^ cells following the protocol described in detail in the original publication (*46*) with no major changes. Due to considerable loss of material during the dual crosslinking and washing steps, we performed the MNase digestion with 100U MNase in 500ul of MB1 buffer assuming that we have roughly 5×10^6 cells at this stage (20U per 1×10^6^ cells). After end-repair and adapter ligation, libraries were amplified with 9 cycles of PCR and sequenced on an Illumina Nova-seq machine (PE50).

### PRO-seq with spiked-in *Drosophila* S2 cells

PRO-seq from resting and activated B cells was performed as described previously (*56*) with several modifications. To isolate nuclei, murine and *Drosphila* S2 cells were resuspended in cold Buffer IA (160 mM Sucrose, 10 mM Tris-Cl pH 8, 3 mM CaCl_2_, 2 mM MgOAc, 0.5% NP-40, 1 mM DTT added fresh), incubated on ice for 3 min and centrifuged at 700 g for 5 min. The pellet was resuspended in nuclei resuspension buffer NRB (50 mM Tris-Cl pH 8, 40% Glycerol, 5 mM MgCl_2_, 0.1 mM EDTA). For each run-on, 10^7^ nuclei of the sample and 10% *Drosophila* S2 nuclei were combined in a total of 100 µL NRB and incubated at 30°C for 3 min with 100 µL 2x NRO buffer including 5µl of all four 1mM Bio-11-NTPs (Perkin-Elmer). Subsequent steps were performed as described (*56*), except that 3’ and 5’ ligations were performed at 16°C overnight with 3’RNA-linker 5’5Phos/NNNNNNNGAUCGUCGGACUGUAGAACUCUGAAC/3InvdT-3’ and 5’-RNA-linker 5’-CCUUGGCACCCGAGAAUUCCANNNN-3, respectively, and CapClip Pyrophosphatase (Biozym Scientific) was used for 5’ end decapping. RNA was reverse transcribed by SuperScript III RT with RP1 Illumina primer to generate cDNA libraries. Libraries were amplified with barcoding Illumina RPI-x primers and the universal reverse primer RP1 using KAPA HiFi Real-Time PCR Library Amplification Kit. Amplified libraries were subjected to gel electrophoresis on 2.5% low melting agarose gel and amplicons from 150 to 350 bp were extracted from the gel, multiplexed and sequenced on Illumina platform HiSeqV4 SR50. Bioinformatic analyses were performed as described (*56*). Library amplification included the use of random 8-mers to exclude PCR duplicates. Libraries were deduplicated followed by trimming of the 8-mer and alignment of reverse complemented reads to the reference genome. For visualization, the penultimate 3’ nucleotide was plotted to represent the true 3’ end of the nascent RNA. Spike-in normalization using a fixed amount of aligned *Drosophila* reads was applied to the samples when indicated (*57*).

### RT-qPCR and ChIP-seq

RT-qPCR with *Drosophila* S2 cell spike-in was performed as described in our previous study (*37*). ChIP-seq for Rad21 (Abcam) was performed exactly as described (*45*).

### Bioinformatics Tri-C

Raw multiplexed sequencing reads were demultiplexed allowing for one mismatch between true and sequenced barcodes. Technical replicates were combined and processed and aligned to the mm9 genome (UCSC) using the previously published CCseqBasicS5 pipeline (https://github.com/Hughes-Genome-Group/CCseqBasicS) adapted to only retain reads with a MAPQ >= 20 and default command line arguments except for --flashBases 10 --ampliconSize 800 --triC --nla. Contact quantifications were merged for each sample with the same condition and the same capture. Tri-C contact matrices were generated from summed contacts using a modified version of the previously published TriC_matrix_simple_MO.py script (https://github.com/oudelaar/TriC) normalizing contacts by the number of restriction sites in each bin and to a total of 300,000 contacts in each matrix.

Contact matrices for each condition were plotted using the average over all biological replicates. The correlation plots were made with the corrplot R package. Capture-C style profiles (1D histograms) were generated from averaged matrices by summing up off-diagonal contacts for each genomic bin and dividing by two to avoid double counting. Contacts on the diagonal were not counted twice and thus added without division. Utilized scripts and code are available at https://github.com/PavriLab/tric.

### Micro-C

Micro-C data were processed using the *distiller* pipeline (https://github.com/open2c/distiller-nf) written for nextflow (*58*). The sequencing reads were aligned to the mouse reference assembly mm9 using *bwa mem* (<u>Li, 2013:</u> https://arxiv.org/abs/1303.3997). The resulting alignments were parsed into Micro-C pairs and filtered for duplicates using the *pairtools* package (https://github.com/open2c/pairtools). These pairs were further aggregated into contact matrices in the cooler format using the *cooler* package at multiple resolutions (*59*) and normalized using the iterative correction procedure (*60*). The contact matrices were visualized using the HiGlass genome browser (*61*).

## ACKNOWLEDGEMENTS

We are especially grateful to Marieke Oudelaar (Max Planck Institute for Biophysical Chemistry, Göttingen, Germany) for extensive discussions and feedback on all aspects of Tri-C, as well as comments of the manuscript. We also thank Douglas Higgs and Jim Hughes (Weatherall Institute for Molecular Medicine, Oxford, UK) for initial discussions on Tri-C. We thank Tsung-Han Hsieh and Robert Tjian (University of California, Berkeley, USA) for advice on Micro-C. We are grateful to the Vienna Biocenter Core Facilities (VBCF) for next-generation sequencing and the IMP/IMBA core facilities especially the animal house, molecular biology and bio-optics services. This work was funded by Boehringer Ingelheim, The Austrian Industrial Research Promotion Agency (Headquarter Grant FFG-834223), and grants from the Austrian Science Fund to UES (FWF T 795-B30) and RP (FWF P 29163-B26).

## AUTHOR CONTRIBUTIONS

JC performed most of the Tri-C assays and analyzed data with supervision from UES. UES performed Tri-C and PRO-seq and analyzed data. DM and MvdL performed bioinformatic analyses. JF performed Micro-C. JF and MM performed Rad21 ChIP-seq. AG analyzed Micro-C data and provided conceptual inputs and critical feedback. RP conceived the project, analyzed data and wrote the manuscript with extensive and critical inputs from JF, UES and MvdL.

---

**Figure S1:**
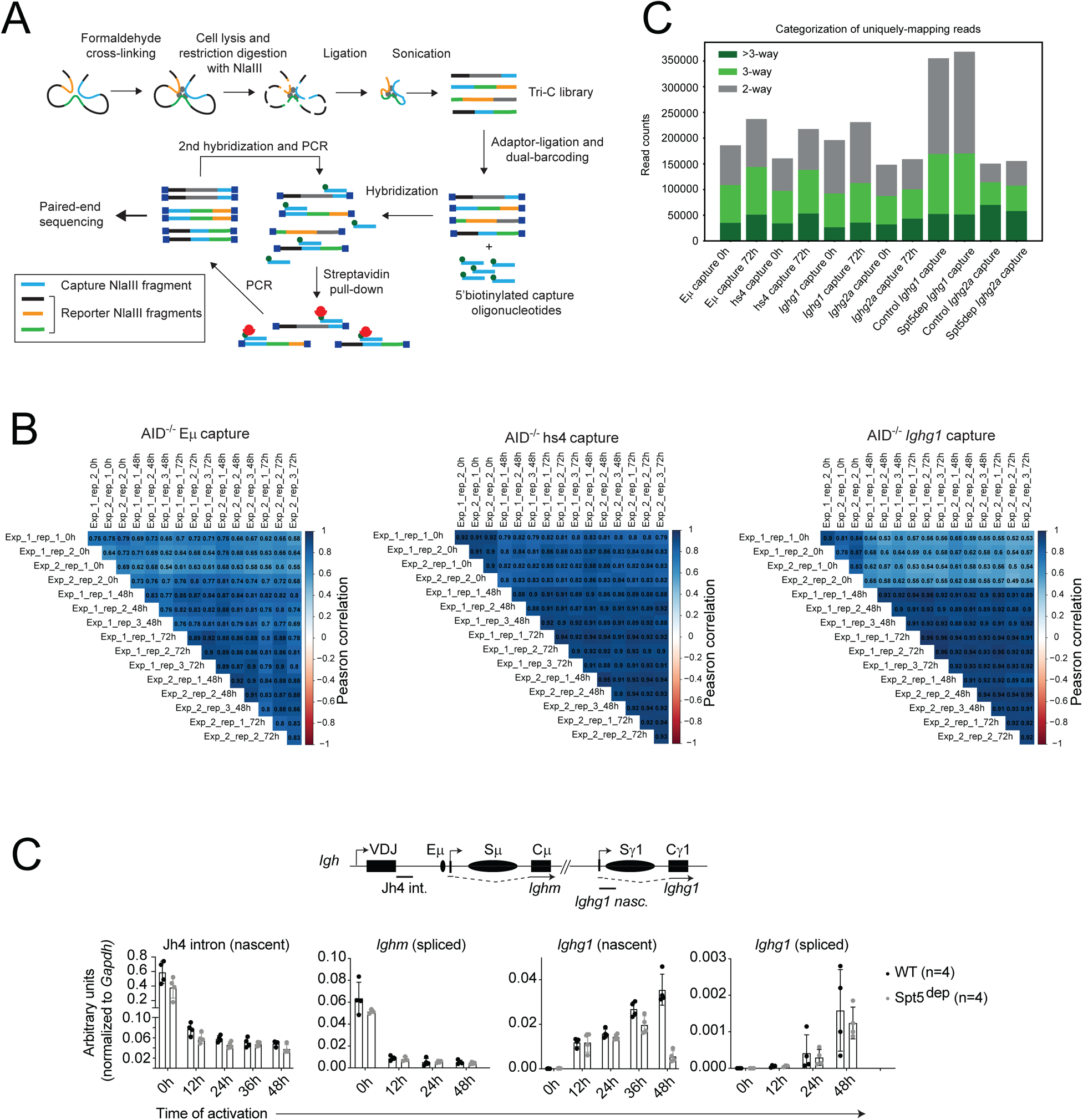
Overview of Tri-C and characterization of Tri-C datasets. A. Overview of Tri-C methodology (adapted from Oudelaar et al. 2018).
B. Correlation between replicates of Tri-C datasets from Eμ, *Ighg1* and hs4 capture viewpoints. The color scale corresponds to the Pearson correlation coefficient shown alongside the matrices.
C. Bar graphs showing the proportion of uniquely-mapping reads containing the capture NlaIII fragment and at least 1 associated reporter NlaIII fragment. Reads are classified as 2-way, 3-way and >3-way if they contain 1, 2 or >2 reporter reads, respectively. The reporter fragments associated with 3-way and >3-way reads are used for generating Tri-C matrices.
D. RT-qPCR analysis of nascent and spliced *Igh* transcripts in WT and Spt5^dep^ cells at the indicated time points post-activation. The qPCR data is the same as in Fig. 3B but was normalized to the levels of the internal, housekeeping *Gapdh* transcript. The location of the transcripts analyzed is shown in the diagram above the graphs.

